# EasyEyes — Accurate fixation for online vision testing of crowding and beyond

**DOI:** 10.1101/2023.07.14.549019

**Authors:** Jan W. Kurzawski, Maria Pombo, Augustin Burchell, Nina M. Hanning, Simon Liao, Najib J. Majaj, Denis G. Pelli

**Affiliations:** New York University

**Author notes:** These authors contributed equally to this work and share first authorship (Jan W. Kurzawski, Maria Pombo).

**Keywords:** fixation, remote testing, online testing, eye-tracker, crowding, EasyEyes, crosshair tracking, gaze control, peripheral testing

## Abstract

Online methods allow testing of larger, more diverse populations, with much less effort than in-lab testing. However, many psychophysical measurements, including visual crowding, require accurate eye fixation, which is classically achieved by testing only experienced observers who have learned to fixate reliably, or by using a gaze tracker to restrict testing to moments when fixation is accurate. Alas, both approaches are impractical online since online observers tend to be inexperienced, and online gaze tracking, using the built-in webcam, has a low precision (±4 deg, Papoutsaki et al., 2016). The EasyEyes open-source software reliably measures peripheral thresholds online with accurate fixation achieved in a novel way, without gaze tracking. EasyEyes tells observers to use the cursor to track a moving crosshair. At a random time during successful tracking, a brief target is presented in the periphery. The observer responds by identifying the target. To evaluate EasyEyes fixation accuracy and thresholds, we tested 12 naive observers in three ways in a counterbalanced order: first, in the lab, using gaze-contingent stimulus presentation (Kurzawski et al., 2023; Pelli et al., 2016); second, in the lab, using EasyEyes while independently monitoring gaze; third, online at home, using EasyEyes. We find that crowding thresholds are consistent (no significant differences in mean and variance of thresholds across ways) and individual differences are conserved. The small root mean square (RMS) fixation error (0.6 deg) during target presentation eliminates the need for gaze tracking. Thus, EasyEyes enables fixation-dependent measurements online, for easy testing of larger and more diverse populations.

## Introduction

Online data collection offers researchers immediate access to thousands of participants around the world, which speeds up research and allows for more diverse samples (Grootswagers, 2020; Palan & Schitter, 2018). However, for researchers conducting visual fixation-dependent experiments, the appeal of online testing is frustrated by the inability to track gaze precisely. This is especially important when stimuli are presented in the periphery.

In peripheral testing, observers are torn between fixating on the central crosshair and looking toward the anticipated target location, which we call “peeking” (Kurzawski et al., 2023). If observers fixate on the anticipated location of the peripheral target, target eccentricity is almost zero, defeating the purpose of peripheral testing. In-lab eye tracking is widely used to ensure fixation. Typically, infrared light emitted by the eye tracker creates a reflection in the cornea of the eye, which is picked up by an infrared camera (*Eye Tracking 101*, 2022). Thus, eye trackers can precisely report the eye’s location and movement at any point in time. However, precise eye trackers are generally expensive, cumbersome, and require calibration. Importantly, they limit the study of fixation-dependent experiments to lab settings.

Precise gaze control is not available for online testing yet. Even though many researchers have devised tools and methods to gather eye-tracking data using participants’ webcams, (Huang et al., 2016; Valliappan et al., 2020; Xu et al., 2015) many of these tools still require calibration. An exception is *WebGazer.js,* (Papoutsaki et al., 2016) a prominent auto-calibrated eye-tracking tool that relies on the webcam to estimate the participant’s gaze. Researchers have shown its effectiveness for various tasks (Semmelmann & Weigelt, 2018; Slim & Hartsuiker, 2022). Nevertheless, in the best-case scenario, its spatial accuracy is about 4 deg, which would introduce a ± 40% error in the eccentricity of a target at 10 deg eccentricity (Huang et al., 2016; Papoutsaki, 2015).

Ample research on eccentricity-based and polar-angle-based differences in perception (see Himmelberg et al., 2023; Strasburger et al., 2011 for reviews) relies on stable central eye fixation (Guzman-Martinez et al., 2009). Visual crowding experiments are well-known fixation-dependent psychophysical tasks. Crowding, or the failure to recognize an object due to clutter, is typically measured by asking participants to recognize a letter between two flankers (Bouma, 1973; Pelli et al., 2004; Pelli & Tillman, 2008; Strasburger, 2020; Stuart & Burian, 1962). *Crowding distance* (“critical spacing”) is the center-to-center distance from target to flanker that achieves a criterion level of performance. It increases with eccentricity, and thus crowding is generally measured in the periphery (Bouma, 1970; Kooi et al., 1994; Levi & Carney, 2009; Pelli et al., 2004; Toet & Levi, 1992).

Crowding varies two-fold across observers (Kurzawski et al., 2023; Pelli et al., 2016) and little within an observer (Chung, 2007; Kurzawski et al., 2023). Clinically, it plays a key role in amblyopia (Levi et al., 2007) and exacerbates the effects of macular degeneration (Wallace et al., 2017). It correlates with dyslexia and thus may be a valuable biomarker to guide early interventions designed to diminish problems in decoding letters and words (Joo et al., 2018; Levi, 2008; Li et al., 2020). For crowding to fulfill its promise as a biomarker, accurate target eccentricity when testing is required. We have previously shown that measured crowding distance depends on fixation accuracy and inaccurate fixation impacts the mean and standard deviation of measured crowding thresholds (Kurzawski et al., 2023). One way to avoid inaccurate fixation is gaze-contingent stimulus presentation that here we call “awaited fixation”: While monitoring gaze with an eye tracker, the stimulus only appears after the observer has accurately fixated for 250 ms. Unfortunately, online gaze tracking is not accurate enough to use this method.

Here we demonstrate how EasyEyes (https://easyeyes.app/), an open-source online psychophysical testing tool, measures crowding thresholds reliably by achieving accurate fixation with a fine motor task and without eye tracking.

Researchers have shown that cursor movement generally correlates with eye movement (Chen et al., 2001; Liebling & Dumais, 2014). Moreover, looking at a target is required for precise and accurate hand movements (Jana et al., 2017), and fixations are necessary when coordinating the movement between two objects (Land & Hayhoe, 2001). With EasyEyes observers perform the fine motor task of tracking a moving crosshair with their cursor. The peripheral target is presented after successful crosshair tracking for a random time of 0.75 to 1.25 sec. This eye-hand coordination task demands accurate fixation before target onset.

We compare thresholds between an at-home online crowding task (EasyEyes home), an in-lab version of the same online task (EasyEyes lab), and a previously validated crowding in-lab task (CriticalSpacing.m lab, Pelli et al., 2016). We find that online EasyEyes crowding thresholds do not significantly differ from those measured in the lab. Additionally, we use gaze tracking while observers complete EasyEyes in the lab to validate that observers fixate on the moving crosshair during target presentation and do not peek.

## Methods

### Observers

12 observers took part in our experiment. Seven identified as female and five as male. Their ages ranged from 21 to 46 (*M* = 27.3, *SD* = 6.8). All observers were fluent English speakers and had normal or corrected-to-normal vision. Importantly, observers were recruited via a convenience sample and ensuring that they had little to no experience with crowding tasks. Two-thirds of the observers were associated with the psychology department of New York University (graduate students, postdocs, and staff), but had no experience with vision psychophysical tasks. All observers gave informed consent in accordance with the Declaration of Helsinki and were compensated $15/hr for their participation. This experiment was approved by the New York University Committee on Activities Involving Human Subjects (UCAIHS; IRB-FY2016-404).

All observers completed a visual crowding task in three ways: 1. CriticalSpacing.m (in lab), 2. EasyEyes (in lab), and 3. EasyEyes (at home). The order of the three ways was counterbalanced across observers to cancel out any order effects.

#### Way 1: CriticalSpacing.m in lab

CriticalSpacing.m (Pelli et al., 2016) is a thoroughly tested MATLAB program for measuring crowding thresholds, recently enhanced by the addition of a chin rest and gaze-contingent display (Kurzawski et al., 2023). The target (with flankers) is presented when gaze is detected within 1.5 deg of the crosshair by an EyeLink 1000 eye tracker. Trials are retained only if fixation remains within 1.5 deg of the crosshair throughout the target duration. Using this method, Kurzawski et al. (2023) report extensive crowding measurements on 50 observers.

#### Way 2: EasyEyes in lab

Observers used the same chin rest, and the EasyEyes online software measured crowding threshold while the EyeLink 1000 independently monitored gaze. The EasyEyes software had no access to the gaze data. An EasyEyes log recorded a time stamp in absolute POSIX time (in fractions of a second), and the crosshair, cursor, and target position every frame (60 Hz). A MATLAB program running in parallel saved a POSIX timestamp and gaze position every 10 ms.

#### Way 3: EasyEyes at home

Each observer opened the URL of the EasyEyes experiment in a browser on their own computer and ran the experiment online.

### Identification task

In all testing methods, observers completed a simple letter recognition task that measures crowding in the visual periphery. In each trial, the observer is presented with a trigram of letters for 150 ms. We refer to the middle letter as the target and the other two as the flankers. For each trial, target and flankers are drawn randomly, without replacement, from a nine-letter character set: DHKNORSVZ. Letters are rendered in black in the Sloan font on a uniform white background of about 275 cd/m^2^ (Pelli et al., 2016; Sloan et al., 1952). We omit the C, from Louise Sloan’s original ten letters, because it is too easily confused with the O (Elliott et al., 1990). The Sloan letters all have the same square (invisible) bounding box. The target letter is presented so that its center is either −10 deg or +10 deg from the fixation crosshair. The flankers are presented, symmetrically, to the right and left of the target. The spacing, center of target to center of each flanker varies from trial to trial, guided by QUEST. After the brief presentation, the list of nine possible letters is displayed, and the observer is asked to identify the target by clicking (or typing, in the case of the EasyEyes home session) one of the nine letters displayed.

Since our observers were naive to the task, they completed a brief (2-3 min) online training session which consisted of 10 trials, 5 at each of ± 10 deg of eccentricity, prior to any session.

### Measuring threshold

In each block, we use QUEST (Watson & Pelli, 1983) to control the letter spacing of each trial and finally estimate the crowding distance threshold. Each threshold estimate was based on 35 trials. Each block of trials interleaved two conditions, one for −10 deg and another for +10 deg (resulting in 35 trials per condition, 70 trials per block). Each participant completed two blocks in each session (140 trials per session). Letter size scales with spacing, maintaining a fixed ratio of 1.4:1. Threshold was defined as the letter spacing for 70% correct identification.

The conditions for target presentation differ depending on the experimental software. CriticalSpacing.m uses gaze-contingent stimulus presentation while EasyEyes relies on crosshair tracking. These are described below.

### Gaze-contingent CriticalSpacing.m

We measured crowding thresholds using CriticalSpacing.m (Pelli et al., 2016) with additional features that integrated compatibility with the EyeLink eye tracker. This enhanced CriticalSpacing.m uses gaze-contingent stimulus presentation that we call “awaited fixation” (Kurzawski et al., 2023). At the start of the experiment, a central crosshair is shown on the screen. The first trial begins when the observer presses the spacebar. After correct fixation for 250 ms, a letter trigram is displayed for 150 ms. After stimulus offset, the observer uses a mouse to click to report the middle letter of the trigram. All possible letters appear in a row below fixation. After clicking on the letter, the observers are instructed to look back at the central crosshair. A correct response is acknowledged with a short beep. Subsequently, the computer waits for the observer to maintain fixation within 1.5 degrees of the crosshair for 250 ms. If the waiting period exceeds 10 sec, the software prompts for recalibration of the gaze tracker.

### Apparatus

In the lab, observers used a chin rest to maintain a 40 cm viewing distance from eye to display. To track gaze in the lab, we used an Eyelink 1000 eye tracker (SR Research, Ottawa, Ontario, Canada) with a sampling rate of 1000 Hz. To allow for a short viewing distance, we used their Tower mount setup with a 25 mm lens.

Each in-lab session was completed with an Apple iMac 27” with an external monitor for stimulus presentation. The screen resolution was 5120 × 2880. Apple iMac has AMD graphics for optimal compatibility with the Psychtoolbox imaging software. The Systems Preference: Displays: Brightness slider was set (by calling MacDisplaySettings.m in the Psychtoolbox) to 0.86 (range 0 to 1) to achieve a white background luminance of about 275 cd/m^2^. The observer viewed the screen binocularly. Stimuli were rendered using *CriticalSpacing*.*m* software (Pelli et al., 2016) implemented in MATLAB 2021 using the Psychtoolbox (Brainard, 1997).

### EasyEyes

EasyEyes (https://easyeyes.app/) is open-access software to measure thresholds online. With a Pavlovia (https://pavlovia.org/) account, the scientist can upload an experiment table with an alphabetical list of parameters along with corresponding files (consent forms and fonts) to the EasyEyes website and obtain an experiment link. EasyEyes integrates Prolific (https://www.prolific.co/) to allow scientists to easily recruit paid participants from all over the world. After participants complete the experiment, EasyEyes provides easy access to the data as well as tools for data analysis and visualization.

EasyEyes has 305 parameters that allow scientists flexibility to include questionnaires and measure various parameters, including reading speed and accuracy, visual acuity, and hearing audiogram. EasyEyes uses the “virtual chinrest” method of Li et al. (2020) to measure screen size and viewing distance and uses Google FaceMesh (Kartynnik et al., 2019) to continuously track viewing distance throughout the experiment.

#### Experimental design

For the EasyEyes version of the letter identification task, we implement the CriticalSpacing.m task described above as closely as possible. Our spreadsheet specifies 3 blocks of two target tasks: one *questionAndAnswer* block (that asks observers for their participant ID and age) and two *identify* blocks. Each *identify* block has two interleaved conditions of 35 trials each. The only difference between the conditions is whether the target position is specified at ±10 degrees (*targetEccentricityXDeg* is 10 or −10 and *targetEccentricityYDeg* is 0). In this way, each block calculates two thresholds, one for the right and one for the left meridian. We specify the threshold criterion proportion correct (70%), the viewing distance (40 cm), and the stimulus presentation time (0.15 s) using the *thresholdProportionCorrect*, *viewingDistanceDesiredCm*, and *targetDurationSec* parameters respectively. We specify *targetKind* to be “letter,” spacingDirection to be “radial,” *spacingRelationToSize* to be “ratio” and *spacingSymmetry* to be “screen”. We also provide the software with the WOFF2 file of the Sloan font and indicate it as such using the *font*, *fontSource*, and *fontCharacterSet* parameters.

There are three differences between the at-home and in-lab EasyEyes experiments. First, the in-lab version sets the *_trackGazeExternallyBool* parameter to TRUE to save a log of timestamped screen locations of crosshair, cursor, and (when present) target. Second, the at-home experiment requires observers to calibrate their screen size and viewing distance as described above. Lastly, in the at-home experiment, the *viewingDistanceNudgingBool* parameter is set to TRUE so observers are told to move farther away from or closer to the screen if their viewing distance is less than 80% or greater than 120% of the specified 40 cm.

#### Moving crosshair

Traditionally, many vision experiments ask observers to fix their gaze on a static fixation mark, which is often a crosshair. Naive observers struggle to hold their gaze on a central mark while anticipating a peripheral target. Instead, EasyEyes tells the observer to use the cursor to track a moving crosshair.

Each trial presents a moving black crosshair consisting of a vertical and a horizontal line crossing at their midpoints, each 2 deg long and 0.05 deg thick. The crosshair has an invisible “hotspot” disk with a 0.1 deg radius about its center location. Until stimulus presentation, the crosshair moves steadily, counterclockwise, along an invisible circular trajectory centered on the screen center, with a radius of 0.5 deg and a period of 10 s, resulting in a speed of 0.3 deg/s. The initial position of the crosshair is a random point on the circular path. The observer is told to use the cursor to track the center of the crosshair. Tracking is considered successful while the cursor tip is in the hotspot, and the crosshair becomes “bold” (increasing line thickness from 0.05 to 0.07 deg) while tracking is successful. This feedback helps the participant to quickly learn tracking.

#### Coordinates

This paper uses two spatial coordinate systems to specify stimulus and gaze position. *Screen coordinates* (pix) are X and Y pixels, with the origin in the upper left corner of the screen (and y increases down). *Visual coordinates* (deg) are X and Y gaze positions after relative to the current location of the crosshair (and y increases up). The target is presented at 10 deg left or right of the crosshair center. (Stimulus display software needs to convert back and forth between these coordinate systems. The supplement provides these routines in MATLAB and JavaScript.)

#### Pre-stimulus interval

At the start of each trial, EasyEyes displays the instruction in the upper left corner of the screen, “Ready? Track the center of the crosshair until the letter(s) appear.” Meanwhile, EasyEyes displays the cursor and moving crosshair, and checks the cursor position every frame until the cursor tip is in the crosshair hotspot, then it starts a timer, which waits for a duration randomly sampled from a uniform probability interval 0.75 to 1.25 s. EasyEyes checks the cursor position again at the end of the (unannounced) tracking interval. If the cursor tip is in the hotspot then EasyEyes presents the target. Otherwise EasyEyes restarts the timer, without disturbing the crosshair’s motion. (Since collecting the data presented here, we made the cursor-tracking criterion more stringent. Now the cursor is checked repeatedly during the tracking interval and any exit from the hot spot causes EasyEyes to go back to the start.)

#### Stimulus and response

The target (with flankers) is displayed immediately, for 150 ms. During the target presentation, the crosshair and cursor are hidden. 700 ms after the stimulus offset, the following instructions appear: “Please identify the middle letter by clicking it below or pressing it on the keyboard.”

### Monitoring gaze while testing with EasyEyes

EasyEyes does not have gaze tracking. We use an in-lab EyeLink 1000 eye tracker to assess the accuracy of fixation during crosshair tracking by the cursor. A simple handshake between EasyEyes and MATLAB (controlling the EyeLink 1000) tells MATLAB when the experiment begins and ends. The handshake uses a RESTful API to a Node.js server. EasyEyes sends a start command by making a POST request to the RESTful server. Meanwhile, MATLAB repeatedly sends GET requests to the server until it receives the “start” command from EasyEyes. For each display frame (60 Hz), EasyEyes records the POSIX (absolute time in fractions of a second) timestamp, and the X and Y screen positions of the crosshair, cursor, and target. MATLAB gets X and Y gaze position from the EyeLink every 10 ms and records it along with a POSIX timestamp. This produces two timestamped CSV files, one from MATLAB and one from EasyEyes, which were combined to generate the plots seen here.

### RMSE of gaze and cursor position

We estimated RMSE by calculating the radial distance between either cursor and crosshair positions (tracking error) or gaze and crosshair positions (gaze pursuit error) for each frame of the stimulus presentation. These errors were averaged within and across trials to produce RMS errors per observer. In both cases (tracking and gaze pursuit errors), we report the mean error across observers.

### Correction of eye tracker calibration offset in X-Y gaze position

Eye tracker is calibrated once before the session by asking the observer to fixate in the center and at 5 deg above, below, right and left from the center. We find that the center of the screen is reported with a small consistent offset unique to each observer session. We estimated and removed this offset. Correction was determined independently for each observer session by calculating the mean X and Y offset between crosshair and recorded gaze position across all gaze samples obtained during the 750 ms interval before stimulus onset (75 gaze samples per trial, and 140 trials). This single offset correction was applied to every gaze position of the observer in that session. Across observers, the mean±SD RMS radial offset was 0.64±0.25 deg.

### Statistical analysis of crowding thresholds

Test-retest correlation was assessed between log crowding distances. We also calculate the test-retest variance (which is reported as SD) as the square root of mean variance across observers. To evaluate the difference between methods, we conducted a one-way ANOVA with log crowding distance as the dependent variable and method (CriticalSpacing.m lab, EasyEyes lab, and EasyEyes home) as the independent variable. Furthermore, we evaluate the difference in log crowding threshold variance with pairwise F-tests for equal variance. To assess whether individual differences are conserved across methods, we compute each observer’s geometric mean threshold (4 thresholds: left and right, test and retest) and calculate the Pearson correlation coefficient for all pairs of methods. To assess how well observer differences are conserved we computed Spearman rank order correlations (of geometric mean across left and right) across methods and across test and retest within each method. For example, to compare CriticalSpacing.m lab to EasyEyes home, we correlate the test thresholds in the former to the retest thresholds in the latter, and vice-versa. We also calculate the intraclass correlation coefficient across all three methods. All analyses were conducted in R (version 4.2.3) using R Studio.

## Results

### Crowding thresholds agree across test-retest

We measured radial crowding thresholds in 12 observers on the right and left meridian at 10 deg of eccentricity using three experimental methods (EasyEyes home, EasyEasy lab, and CriticalSpacing.m lab). To assess the reliability of each method, we tested each threshold twice in two blocks separated by a break. We find that for all methods the test-retest correlations are highly significant (p < 0.1). The test-retest standard deviation was similar across methods (**Figure 1A**) and was not different from the results for 50 observers tested by Kurzawski et al. (2023) in an in-lab setting. Summary of standard deviations for our three methods and Kurzawski et al. (2023) are shown in **Table 1**. **Figure 1A** directly compares test and retest thresholds across methods. In each method, crowding distance was around 0.8 smaller in the retest than in the test session, slightly smaller than the 0.9 in previous reports (Chung, 2007; Kurzawski et al., 2023). The improvement of the second session was independent of which method was used first to test the observer. Overall, crowding thresholds based on one 35-trial QUEST staircase have similarly good reproducibility across all three methods.

**Figure 1.**
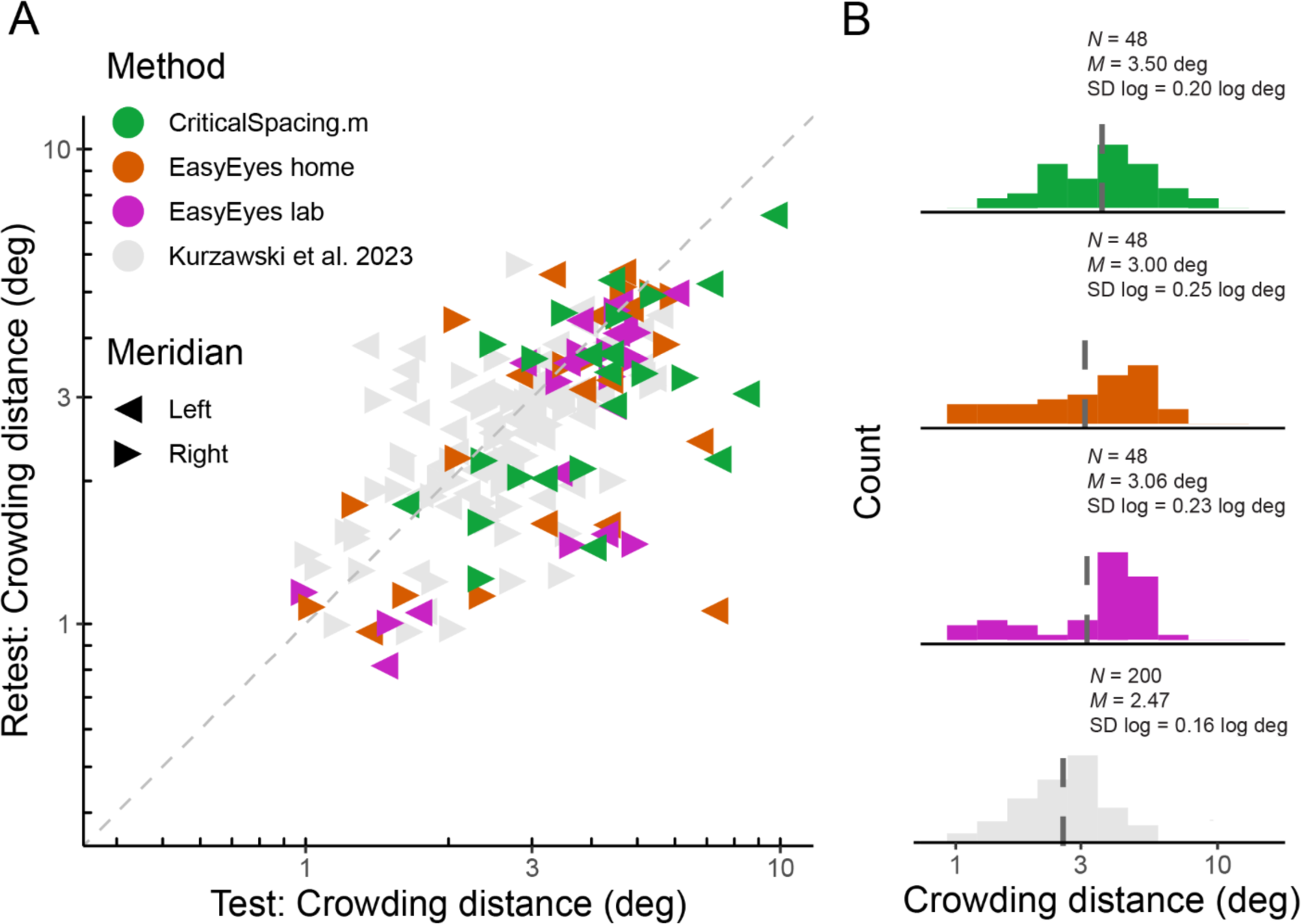
Log crowding thresholds across methods. **A**. Test-retest crowding distances across methods. Gray triangles are thresholds measured by Kurzawski et al. (2023), colored triangles are data acquired for the purpose of this project. Axes are log-log. **B**. Histograms of log thresholds across methods. Dashed lines indicate the geometric mean. *N* indicates number of observers, *M* is a geometric mean and SD is the standard deviation of all measured log crowding distances (12 observers, two meridians, test and retest) for our data and for fraction of data from Kurzawski et al., (50 observers, two meridians, test-retest).

**Table 1.** Comparing test-retest thresholds for all methods. Test-retest SD represents the mean across observers of the standard deviation of test and retest of log crowding distance.

| Method of measuring crowding distance test-retest | Test-retest SD | Pearson's $r$ | Pearson's $p$ | Ratio |
| --- | --- | --- | --- | --- |
| Critical Spacing.m lab | 0.16 | 0.58 | <0.01 | 0.78 |
| Easy Eyes (home) | 0.18 | 0.53 | <0.01 | 0.82 |
| Easy Eyes (lab) | 0.14 | 0.76 | <0.01 | 0.79 |
| Kurzawski et al., (2023) | 0.11 | 0.55 | <0.01 | 0.94 |

### Crowding thresholds agree across methods

Crowding varies across observers and very little across sessions within an observer (Chung, 2007; Kurzawski et al., 2023). While it is important to assess crowding’s reproducibility within a testing session or experiment, the core of this study is to compare crowding thresholds across three methods. Despite the variations between these methods, we find no significant differences between their measured thresholds. A one-way ANOVA shows no significant difference in mean log crowding threshold estimates, *F*(2) = 1.19, *p* = 0.308. Pairwise F-tests of equal variance show no significant difference in the log variance across methods: *F*(47, 47) = 1.26, *p* = 0.429 (Easy Eyes lab vs. CriticalSpacing.m lab), *F*(47, 47) = 1.49, *p* = 0.173 (Easy Eyes home vs. CriticalSpacing.m lab), and *F*(47, 47) = 0.85, *p* = 0.566 (Easy Eyes lab vs. Easy Eyes home).

Geometric mean (and SD log) was 3.06 (0.23) deg for EasyEyes lab, 3.00 (0.25) deg for Easyeyes home, and 3.5 (0.20) deg for CriticalSpacing.m lab (**Figure 1B**). Additionally, these estimates closely resembled the 50-observer crowding survey published by Kurzawski et al. (2023) 2.47 (0.16) deg.

### Individual differences are conserved across methods

Here we check whether individual differences are reproducible across the three methods. Pearson’s correlation coefficients across methods are high, showing that these differences were conserved (**Figure 2A**). Furthermore, the test-retest correlations within each method are not different from test-retest across methods (**Figure 2B**). This is indicated by similar values of Spearman’s rank correlation coefficients across the whole correlation matrix. To evaluate the consistency across all three methods, we calculated the intraclass correlation coefficient (ICC), which was 0.77 and indicates good reliability (Koo & Li, 2016).

**Figure 2.**
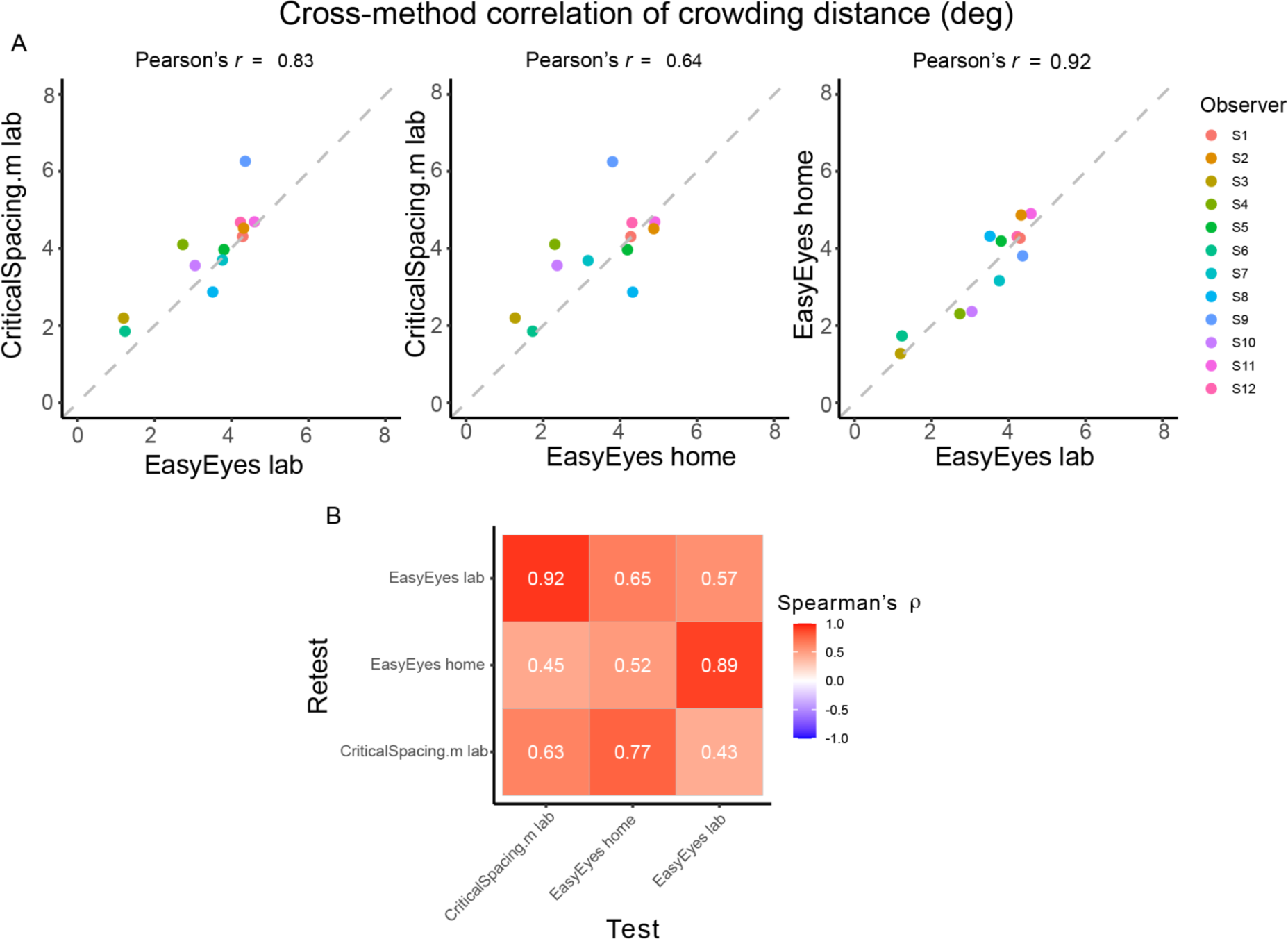
Correlations of crowding distance across methods. **A**. The cross-method correlations of the geometric mean crowding distance for each observer. Mean is calculated from 4 thresholds (2 meridians, test-retest). **B**. Test-retest Spearman’s correlation across mean crowding distance thresholds, across and within methods.

### Fixational accuracy of EasyEyes

Observers are asked to use the cursor to track the moving crosshair, which they do quite well (RMSE of 0.08 deg). During tracking, the target appears at a random time (between 0.75 - 1.25 s), so the observer cannot predict when to look toward the anticipated target location. When the timed interval ends, the crosshair disappears, the cursor is hidden, and the target appears.

EasyEyes does not monitor gaze. We used gaze tracking to monitor how well this foveal tracking task achieves correct fixation during stimulus presentation. For reference, we similarly analyze the conventional awaited-fixation method (gaze-contingent stimulus presentation), which uses gaze tracking (Hanning et al., 2022; Hanning & Deubel, 2022; Kreyenmeier et al., 2020; Kurzawski et al., 2023; Kwak et al., 2023).

While observers are tracking, their gaze remains near the crosshair (RMS of 0.6 deg). **Figure 3A** shows X and Y screen coordinates of gaze, cursor, and crosshair position as a function of time relative to stimulus onset. **Figure 3B** traces the X and Y position of gaze, crosshair, and cursor for one trial per participant during the last 750 ms of tracking.

**Figure 3.**
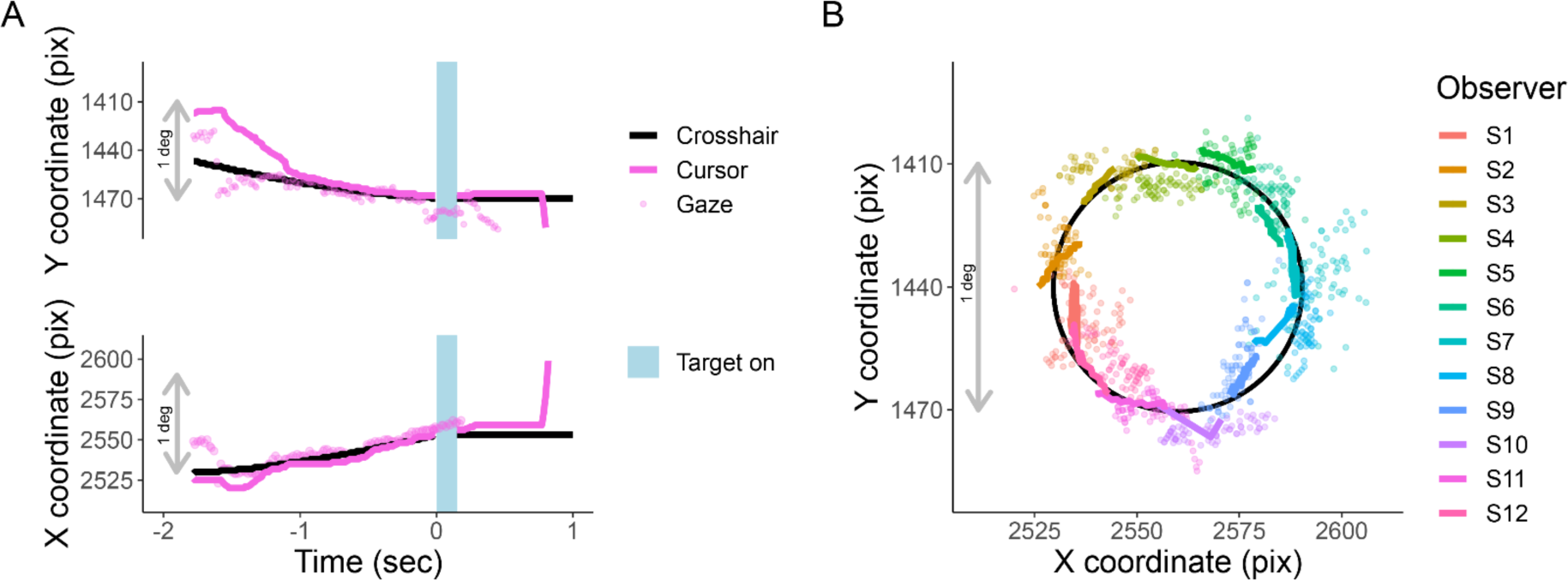
Cursor tracking task. X and Y positions of crosshair (solid black line), cursor (solid colored line), and gaze (colored points) during an EasyEyes trial. The gray bar corresponds to 1 degree (60 pix/deg). **A**. X and Y coordinates as a function of time relative to stimulus onset of one observer. The light blue bar represents the target duration (150ms). **B**. Shows a single representative trial from each observer (750 ms before target onset). The black circle is the trajectory of the crosshair. Again, thick colored lines indicate cursor, and colored dots indicate gaze position. Each observer’s data has been rotated around a circle (crosshair’s trajectory) to minimize overlap with other observers. The pink trial in panel A corresponds to S12 plotted in panel B. All X and Y positions have been corrected for estimated calibration bias.

### Reliability of fixation with EasyEyes

**Figure 3B** presented single-trial gaze tracking before stimulus presentation. **Figure 4** shows gaze (visual coordinates), before, during, and after the stimulus presentation. Visual coordinates are in degrees relative to the center of the crosshair. The mean gaze position (obtained every 10 ms) across all trials (140), and all 12 observers is within 0.03 deg of the crosshair before and during the stimulus presentation, and within 0.3 deg after stimulus presentation. The standard deviation is the lowest in the pre-stimulus interval, while observers are tracking (SD X = 0.97 deg, SD Y = 0.48 deg), increases during stimulus presentation (SD X = 2.03 deg, SD Y = 0.9 deg) and is highest after stimulus offset (SD X = 6.36 deg, SD Y = 1.67 deg). The higher standard deviation in horizontal direction before and during stimulus presentation reflects the overall tendency of eye movements to be directed horizontally (Engbert, 2006; Najemnik & Geisler, 2008; Otero-Millan et al., 2013; Rolfs & Schweitzer, 2022). The pronounced variance in horizontal gaze position after stimulus offset indicates (saccadic) eye movements toward the (now absent) target – a phenomenon commonly referred to as “looking at nothing” (Ferreira et al., 2008). Note that this gaze variance does not affect our perceptual measurement, as the target was already undrawn. For gaze deviations during stimulus presentation see **Table 2**.

**Figure 4.**
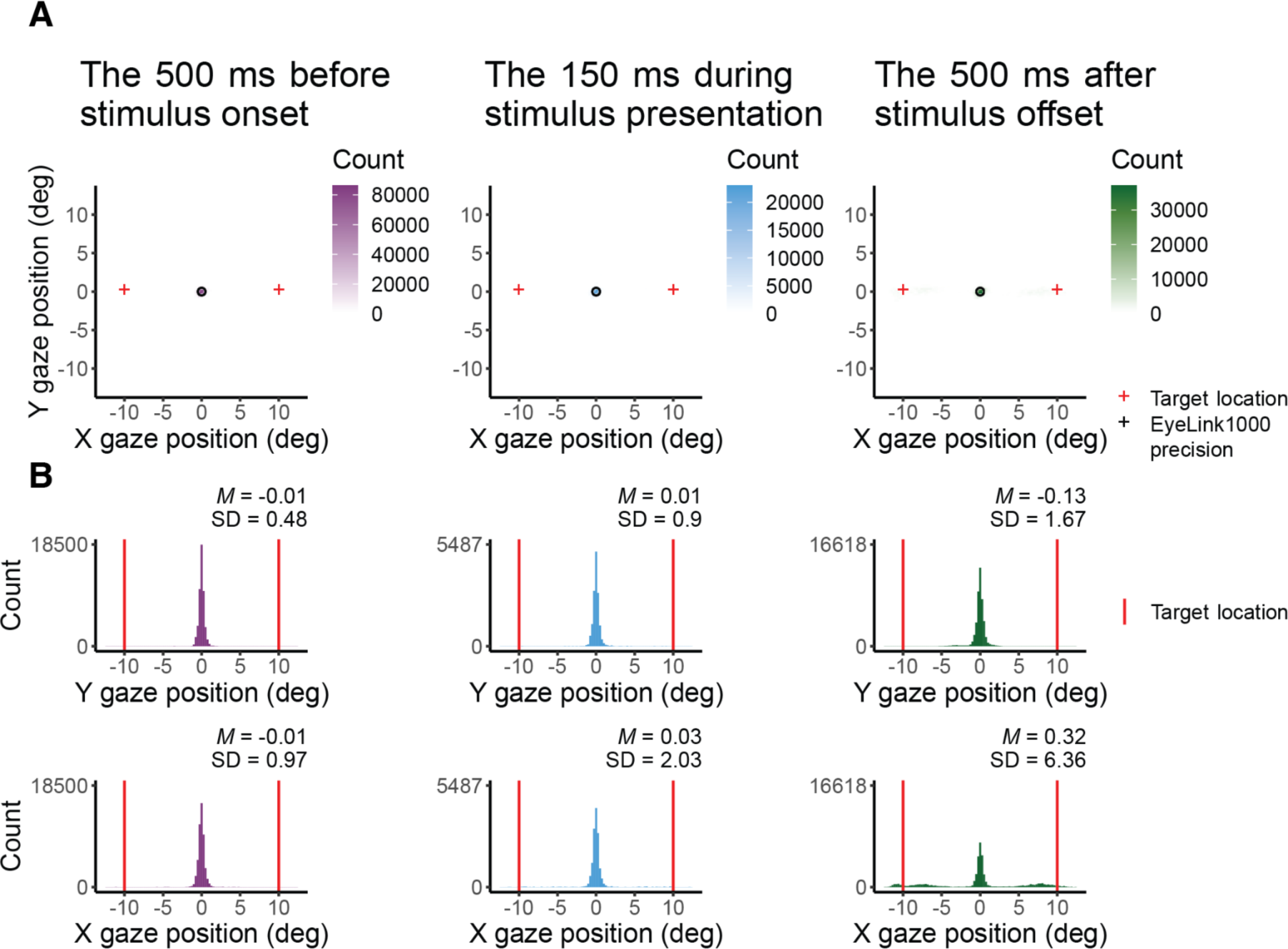
X and Y coordinates (in degrees) of gaze before, during, and after target presentation. **A** shows 2D histograms of gaze. The circle indicates the eye tracker precision and the red cross the target location. **B** shows gaze position in X and Y coordinates. The red vertical line indicates the target location.

**Table 2.** Standard deviations (in deg) of gaze position in X and Y coordinates during stimulus presentation for each participant using EasyEyes lab.

| Observer | SD X (deg) | SD Y (deg) |
| --- | --- | --- |
| S1 | 0.58 | 1.43 |
| S2 | 0.45 | 0.37 |
| S3 | 1.84 | 1.38 |
| S4 | 1.02 | 0.93 |
| S5 | 1.51 | 0.43 |
| S6 | 0.31 | 0.16 |
| S7 | 0.62 | 0.48 |
| S8 | 0.45 | 0.52 |
| S9 | 6.54 | 1.92 |
| S10 | 0.53 | 0.34 |
| S11 | 0.68 | 0.50 |
| S12 | 0.37 | 0.25 |

One out of 12 observers (S9) has a much higher standard deviation (**Table 2**) and often peeked at around 120 ms after stimulus onset. The short latency indicates that this participant has planned the eye movements with target onset. The same observer also peeked during many CriticalSpacing.m lab trials where they were detected and rejected by gaze tracking (**Figure 5**). The eleven remaining observers showed negligible peeking.

**Figure 5.**
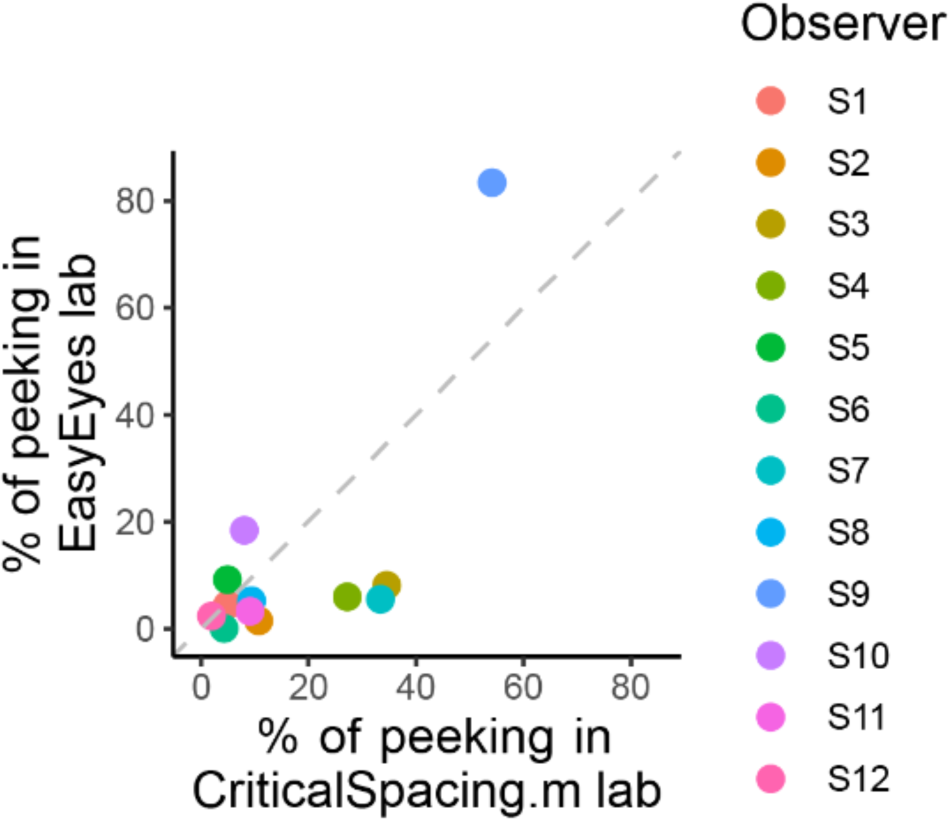
Comparing peeking across methods. The plot shows the percentage of trials in which observers peeked, that is their gaze position was more than 1.5 deg away from the crosshair during stimulus presentation. For CriticalSpacing.m lab peeks are detected by the eye tracker and correspond to rejected trials. For EasyEyes lab we use gaze data to calculate the percentage of peeks post-hoc.

### Less peeking with moving crosshair task than with static fixation

Criticalspacing.m lab counts the peeks and repeats the trials in which observers peeked.

While using EasyEyes lab we monitored gaze position to count each observer’s peeks. In the last section we showed that observers fixate near the crosshair while tracking it with a cursor. Here, we wondered if the tracking also reduces the urge to peek. We compared how often observers peek using a stationary crosshair method in CriticalSpacing.m lab versus the moving crosshair tracking task with EasyEyes lab. A peek is deemed to occur when the observer’s gaze deviates more than 1.5 deg from the last crosshair center position during stimulus presentation. The tracking task roughly halved the number of peeks across observers (median decreased from 9.3% to 5.4%). Individual data is shown in **Figure 5**. Even with a rather short stimulus duration (150 ms) one observer managed to peek with both methods. However their peeking did not change their mean crowding distance relative to other observers. Furthermore their thresholds were consistent across methods even though CriticalSpacing.m lab rejects peeking trial and EasyEyes cannot.

## Discussion

EasyEyes offers a new task to achieve accurate fixation online. We evaluated the accuracy of fixation and compared crowding thresholds measured online with EasyEyes or with our in-lab method (Kurzawski et al., 2023).

We tested 12 naive observers using traditional fixation and gaze-contingent stimulus display (CriticalSpacing.m lab), and using EasyEyes online at home (EasyEyes home) as well as in the lab while independently monitoring gaze (EasyEyes lab). Comparing the mean and standard deviation in thresholds across observers, we do not find significant differences across methods. Cross-method and within-method correlations are not different and individual differences are conserved. With a gaze tracker, we validate that EasyEyes achieves accurate fixation during target presentation.

### Importance of accurate fixation

Visual sensitivity decays with increasing distance from the fovea (the center of gaze). The density of photoreceptor and midget retinal ganglion cells declines with retinal eccentricity, increasing receptive field size (Anton-Erxleben & Carrasco, 2013; Freeman & Simoncelli, 2011). Thus, the visual system loses sensitivity (e.g., to higher spatial frequencies, contrast, or orientation changes) in the periphery. As a consequence, performance in nearly all visual tasks scales with the retinal eccentricity of the test stimulus (but see Hanning & Deubel, 2022 for an eccentricity-independent approach). Visual crowding is no exception: crowding distance scales linearly with eccentricity (Bouma, 1970). In order to achieve a stable threshold estimate, precise control of fixation is indispensable to ensure consistent measurement at the desired retinal eccentricity.

### Gaze during cursor tracking

Previous research has shown that successful tracking of a moving object with a hand-controlled cursor requires that gaze closely follow the moving object (Danion & Flanagan, 2018; Niehorster et al., 2015; Xia & Barnes, 1999). Based on this, we asked observers to use the cursor to track the moving crosshair with the goal of keeping their gaze near the crosshair. Indeed, all of our observers use the cursor reliably to track the crosshair, keeping their gaze near both. This ensures the desired retinal target eccentricity. Both gaze and hand tend to lag the target (Koken & Erkelens, 1992).

### Comparing to Kurzawski et al 2023

While our study is very similar to Kurzawski et al. (2023) –both used CriticalSpacing.m– we find several minor differences compared to their thresholds. The test-retest standard deviation was slightly lower in Kurzawski et al. (2023) than the one reported here (0.11 versus 0.16). The 50 participants in Kurzawski et al. were graduate psychology students with experience in peripheral testing, while here we recruited 12 adults (mean age 27 years) in the university area with no experience in psychophysical testing. We hypothesize that data obtained from experienced observers is less noisy and results are thus more consistent. Crowding is subject to training effects. While observers practice and become more familiar with the task, their crowding distance decreases (Chung, 2007). We find that the geometric mean of crowding distance in Kurzawski et al. (2023) was 2.5 deg while here it was 3.5 deg, i.e., the more experienced observers had lower thresholds. Despite these differences, the consistency across methods here is high, and individual differences in crowding distance are conserved.

### Why measure crowding (online)?

Both crowding distance and acuity are roughly proportional to eccentricity (Bouma, 1970) and thus are similarly sensitive to errors in fixation. We are not aware of any test that is more sensitive to eccentricity. Ophthalmology and optometry clinics routinely measure acuity. Here we explore the possibility that they might find it worthwhile to also measure crowding. Foveal acuity determines the smallest text size that can be read at a certain eccentricity, and peripheral crowding puts an upper limit on reading speed (Pelli et al., 2007). Kurzawski et al., (2023) found hardly any correlation (*r* = 0.15) between foveal and peripheral crowding. Because of its sensitivity to eccentricity and its potential clinical utility, peripheral crowding is a suitable measurement to validate EasyEyes.

From a scientific point of view, accurate fixation for online vision testing enabled by EasyEyes will help to scale up our study of crowding as a promising biomarker of development and health of the visual cortex. Crowding is correlated with dyslexia (Kwon et al., 2007), and can be measured years before the child learns to read (Pelli & Tillman, 2008). Besides this, online testing will facilitate cross-sectional and longitudinal surveys of crowding and related measures. Based on its correlation with dyslexia, we also anticipate a correlation between crowding and reading speed, and that pre-literate crowding might predict later reading speed.

### Why test online?

Online testing allows researchers to test hundreds or thousands of participants in a day, recruit diverse and special populations, and screen underserved populations. As online vision testing gains popularity, a new generation of testing software (e.g., jsPsych (de Leeuw, 2015), lab.js (Henninger et al., 2019), PsychoJS (Pitiot et al., 2017), Gorilla (Anwyl-Irvine et al., 2020), OpenSesame (Mathôt et al., 2012)) makes it easier to test online than in the lab. However, established software lacks the possibility of using gaze tracking to achieve precise fixation – a requirement for most vision tests. Using the cursor to track a moving crosshair, EasyEyes delivers precise fixation – and the same thresholds online as we previously have measured in the lab. Our study shows that EasyEyes is a promising tool for lab-quality online vision testing. Despite other differences, such as absence of supervision, diversity of equipment, and domestic distractions, EasyEyes achieves precise peripheral, fixation-dependent measurements that so far could only be obtained in the lab

## Conclusion

We compare crowding thresholds measured online with EasyEyes or measured in the lab. We find no differences in the mean and standard deviation across an at-home online crowding task (EasyEyes home), an in-lab version of the same online task (EasyEyes lab), and a previously validated crowding in-lab task (CriticalSpacing.m lab). Cursor tracking of a moving crosshair yields accurate fixation (RMSE 0.6 deg) during stimulus presentation. This trick facilitates online testing of peripheral thresholds and any fixation-dependent measures. EasyEyes enables fixation-dependent measurements online, for easy testing of larger and more diverse populations.

## Acknowledgments

This research was funded by the R01-EY031446 to NJM. It was also supported by an NIH core vision grant P30-EY013079 and by a Marie Skłodowska-Curie individual fellowship by the European Commission (898520) to NMH.

## References

Anton-Erxleben, K., & Carrasco, M. (2013). Attentional enhancement of spatial resolution: Linking behavioural and neurophysiological evidence. Nature Reviews. Neuroscience, 14(3), 188–200. https://doi.org/10.1038/nrn3443

Anwyl-Irvine, A. L., Massonnié, J., Flitton, A., Kirkham, N., & Evershed, J. K. (2020). Gorilla in our midst: An online behavioral experiment builder. Behavior Research Methods, 52(1), 388–407. https://doi.org/10.3758/s13428-019-01237-x

Bouma, H. (1970). Interaction Effects in Parafoveal Letter Recognition. Nature, 226(5241), Article 5241. https://doi.org/10.1038/226177a0

Bouma, H. (1973). Visual interference in the parafoveal recognition of initial and final letters of words. Vision Research, 13(4), 767–782. https://doi.org/10.1016/0042-6989(73)90041-2

Brainard, D. H. (1997). The Psychophysics Toolbox. Spatial Vision, 10(4), 433–436. https://doi.org/10.1163/156856897X00357

Chen, M. C., Anderson, J. R., & Sohn, M. H. (2001). What can a mouse cursor tell us more? Correlation of eye/mouse movements on web browsing. CHI ‘01 Extended Abstracts on Human Factors in Computing Systems, 281–282. https://doi.org/10.1145/634067.634234

Chung, S. T. L. (2007). Learning to identify crowded letters: Does it improve reading speed? Vision Research, 47(25), 3150–3159. https://doi.org/10.1016/j.visres.2007.08.017

Danion, F. R., & Flanagan, J. R. (2018). Different gaze strategies during eye versus hand tracking of a moving target. Scientific Reports, 8(1), Article 1. https://doi.org/10.1038/s41598-018-28434-6

de Leeuw, J. R. (2015). jsPsych: A JavaScript library for creating behavioral experiments in a Web browser. Behavior Research Methods, 47(1), 1–12. https://doi.org/10.3758/s13428-014-0458-y

Elliott, D. B., Whitaker, D., & Bonette, L. (1990). Differences in the legibility of letters at contrast threshold using the Pelli-Robson chart. Ophthalmic and Physiological Optics, 10(4), 323–326. https://doi.org/10.1111/j.1475-1313.1990.tb00877.x

Engbert, R. (2006). Microsaccades: A microcosm for research on oculomotor control, attention, and visual perception. Progress in Brain Research, 154, 177–192. https://doi.org/10.1016/S0079-6123(06)54009-9

Eye Tracking 101: What Is It & How Does It Work In Real Life? - Eyeware. (2022, March 3). https://eyeware.tech/blog/what-is-eye-tracking/

Ferreira, F., Apel, J., & Henderson, J. M. (2008). Taking a new look at looking at nothing. Trends in Cognitive Sciences, 12(11), 405–410. https://doi.org/10.1016/j.tics.2008.07.007

Freeman, J., & Simoncelli, E. P. (2011). Metamers of the ventral stream. Nature Neuroscience, 14(9), Article 9. https://doi.org/10.1038/nn.2889

Grootswagers, T. (2020). A primer on running human behavioural experiments online. Behavior Research Methods, 52(6), 2283–2286. https://doi.org/10.3758/s13428-020-01395-3

Guzman-Martinez, E., Leung, P., Franconeri, S., Grabowecky, M., & Suzuki, S. (2009). Rapid eye-fixation training without eye tracking. Psychonomic Bulletin & Review, 16(3), 491–496. https://doi.org/10.3758/PBR.16.3.491

Hanning, N. M., & Deubel, H. (2022). A dynamic 1/f noise protocol to assess visual attention without biasing perceptual processing. Behavior Research Methods. https://doi.org/10.3758/s13428-022-01916-2

Hanning, N. M., Himmelberg, M. M., & Carrasco, M. (2022). Presaccadic attention enhances contrast sensitivity, but not at the upper vertical meridian. IScience, 25(2), 103851. https://doi.org/10.1016/j.isci.2022.103851

Henninger, F., Shevchenko, Y., Mertens, U., Kieslich, P. J., & Hilbig, B. E. (2019). lab.js: A free, open, online experiment builder. Zenodo. https://doi.org/10.5281/zenodo.2775942

Himmelberg, M. M., Winawer, J., & Carrasco, M. (2023). Polar angle asymmetries in visual perception and neural architecture. Trends in Neurosciences, 46(6), 445–458. https://doi.org/10.1016/j.tins.2023.03.006

Huang, M. X., Kwok, T. C. K., Ngai, G., Chan, S. C. F., & Leong, H. V. (2016). Building a Personalized, Auto-Calibrating Eye Tracker from User Interactions. Proceedings of the 2016 CHI Conference on Human Factors in Computing Systems, 5169–5179. https://doi.org/10.1145/2858036.2858404

Jana, S., Gopal, A., & Murthy, A. (2017). A Computational Framework for Understanding Eye–Hand Coordination. Journal of the Indian Institute of Science, 97(4), 543–554. https://doi.org/10.1007/s41745-017-0054-0

Joo, S. J., White, A. L., Strodtman, D. J., & Yeatman, J. D. (2018). Optimizing text for an individual’s visual system: The contribution of visual crowding to reading difficulties. Cortex, 103, 291–301. https://doi.org/10.1016/j.cortex.2018.03.013

Kartynnik, Y., Ablavatski, A., Grishchenko, I., & Grundmann, M. (2019). Real-time Facial Surface Geometry from Monocular Video on Mobile GPUs (arXiv:1907.06724). arXiv. http://arxiv.org/abs/1907.06724

Koken, P. W., & Erkelens, C. J. (1992). Influences of hand movements on eye movements in tracking tasks in man. Experimental Brain Research, 88(3), 657–664. https://doi.org/10.1007/BF00228195

Koo, T. K., & Li, M. Y. (2016). A Guideline of Selecting and Reporting Intraclass Correlation Coefficients for Reliability Research. Journal of Chiropractic Medicine, 15(2), 155–163. https://doi.org/10.1016/j.jcm.2016.02.012

Kooi, F., Toet, A., Tripathy, S., & Levi, D. (1994). The effect of similarity and duration on spatial interaction in peripheral vision. Spatial Vision, 8, 255–279. https://doi.org/10.1163/156856894X00350

Kreyenmeier, P., Deubel, H., & Hanning, N. M. (2020). Theory of visual attention (TVA) in action: Assessing premotor attention in simultaneous eye-hand movements. Cortex: A Journal Devoted to the Study of the Nervous System and Behavior, 133, 133–148. https://doi.org/10.1016/j.cortex.2020.09.020

Kurzawski, J. W., Burchell, A., Thapa, D., Winawer, J., Majaj, N. J., & Pelli, D. G. (2023). The Bouma law accounts for crowding in fifty observers (p. 2021.04.12.439570). bioRxiv. https://doi.org/10.1101/2021.04.12.439570

Kwak, Y., Hanning, N. M., & Carrasco, M. (2023). Presaccadic attention sharpens visual acuity. Scientific Reports, 13(1), Article 1. https://doi.org/10.1038/s41598-023-29990-2

Kwon, M., Legge, G. E., & Dubbels, B. R. (2007). Developmental changes in the visual span for reading. Vision Research, 47(22), 2889–2900. https://doi.org/10.1016/j.visres.2007.08.002

Land, M. F., & Hayhoe, M. (2001). In what ways do eye movements contribute to everyday activities? Vision Research, 41(25), 3559–3565. https://doi.org/10.1016/S0042-6989(01)00102-X

Levi, D. M. (2008). Crowding—An essential bottleneck for object recognition: A mini-review. Vision Research, 48(5), 635–654. https://doi.org/10.1016/j.visres.2007.12.009

Levi, D. M., & Carney, T. (2009). Crowding in Peripheral Vision: Why Bigger Is Better. Current Biology, 19(23), 1988–1993. https://doi.org/10.1016/j.cub.2009.09.056

Levi, D. M., Song, S., & Pelli, D. G. (2007). Amblyopic reading is crowded. Journal of Vision, 7(2), 1–17. https://doi.org/10.1167/7.2.21

Li, Q., Joo, S. J., Yeatman, J. D., & Reinecke, K. (2020). Controlling for Participants’ Viewing Distance in Large-Scale, Psychophysical Online Experiments Using a Virtual Chinrest. Scientific Reports, 10(1), Article 1. https://doi.org/10.1038/s41598-019-57204-1

Liebling, D. J., & Dumais, S. T. (2014). Gaze and mouse coordination in everyday work. Proceedings of the 2014 ACM International Joint Conference on Pervasive and Ubiquitous Computing: Adjunct Publication, 1141–1150. https://doi.org/10.1145/2638728.2641692

Mathôt, S., Schreij, D., & Theeuwes, J. (2012). OpenSesame: An open-source, graphical experiment builder for the social sciences. Behavior Research Methods, 44(2), 314–324. https://doi.org/10.3758/s13428-011-0168-7

Najemnik, J., & Geisler, W. S. (2008). Eye movement statistics in humans are consistent with an optimal search strategy. Journal of Vision, 8(3), 4. https://doi.org/10.1167/8.3.4

Niehorster, D. C., Siu, W. W. F., & Li, L. (2015). Manual tracking enhances smooth pursuit eye movements. Journal of Vision, 15(15), 11. https://doi.org/10.1167/15.15.11

Otero-Millan, J., Macknik, S. L., Langston, R. E., & Martinez-Conde, S. (2013). An oculomotor continuum from exploration to fixation. Proceedings of the National Academy of Sciences of the United States of America, 110(15), 6175–6180. https://doi.org/10.1073/pnas.1222715110

Palan, S., & Schitter, C. (2018). Prolific.ac—A subject pool for online experiments. Journal of Behavioral and Experimental Finance, 17, 22–27. https://doi.org/10.1016/j.jbef.2017.12.004

Papoutsaki, A. (2015). Scalable Webcam Eye Tracking by Learning from User Interactions. Proceedings of the 33rd Annual ACM Conference Extended Abstracts on Human Factors in Computing Systems, 219–222. https://doi.org/10.1145/2702613.2702627

Papoutsaki, A., Sangkloy, P., Laskey, J., Daskalova, N., Huang, J., & Hays, J. (2016). WebGazer: Scalable Webcam Eye Tracking Using User Interactions.

Pelli, D. G., Palomares, M., & Majaj, N. J. (2004). Crowding is unlike ordinary masking: Distinguishing feature integration from detection. Journal of Vision, 4(12), 12–12.

Pelli, D. G., & Tillman, K. A. (2008). The uncrowded window of object recognition. Nature Neuroscience, 11(10), Article 10. https://doi.org/10.1038/nn.2187

Pelli, D. G., Tillman, K. A., Freeman, J., Su, M., Berger, T. D., & Majaj, N. J. (2007). Crowding and eccentricity determine reading rate. Journal of Vision, 7(2), 20. https://doi.org/10.1167/7.2.20

Pelli, D. G., Waugh, S. J., Martelli, M., Crutch, S. J., Primativo, S., Yong, K. X., Rhodes, M., Yee, K., Wu, X., & Famira, H. F. (2016). A clinical test for visual crowding. F1000Research, 5, 81.

Pitiot, A., Agafonov, N., Bakagiannis, S., Pierce, J., Pronk, T., Sogo, H., & Zhao, S. (2017). PsychoJS [JavaScript]. PsychoPy. https://github.com/psychopy/psychojs

Rolfs, M., & Schweitzer, R. (2022). Coupling perception to action through incidental sensory consequences of motor behaviour. Nature Reviews Psychology, 1(2), Article 2. https://doi.org/10.1038/s44159-021-00015-x

Semmelmann, K., & Weigelt, S. (2018). Online webcam-based eye tracking in cognitive science: A first look. Behavior Research Methods, 50(2), 451–465. https://doi.org/10.3758/s13428-017-0913-7

Slim, M. S., & Hartsuiker, R. J. (2022). Moving visual world experiments online? A web-based replication of Dijkgraaf, Hartsuiker, and Duyck (2017) using PCIbex and WebGazer.js. Behavior Research Methods. https://doi.org/10.3758/s13428-022-01989-z

Sloan, L. L., Rowland, W. M., & Altman, A. (1952). Comparison of three types of test target for the measurement of visual acuity. 8(1), 4–16.

Strasburger, H. (2020). Seven Myths on Crowding and Peripheral Vision. I-Perception, 11(3), 2041669520913052. https://doi.org/10.1177/2041669520913052

Strasburger, H., Rentschler, I., & Jüttner, M. (2011). Peripheral vision and pattern recognition: A review. Journal of Vision, 11(5), 13. https://doi.org/10.1167/11.5.13

Stuart, J. A., & Burian, H. M. (1962). A Study of Separation Difficulty*: Its Relationship to Visual Acuity in Normal and Amblyopic Eyes. American Journal of Ophthalmology, 53(3), 471–477. https://doi.org/10.1016/0002-9394(62)94878-X

Toet, A., & Levi, D. M. (1992). The two-dimensional shape of spatial interaction zones in the parafovea. Vision Research, 32(7), 1349–1357. https://doi.org/10.1016/0042-6989(92)90227-A

Valliappan, N., Dai, N., Steinberg, E., He, J., Rogers, K., Ramachandran, V., Xu, P., Shojaeizadeh, M., Guo, L., Kohlhoff, K., & Navalpakkam, V. (2020). Accelerating eye movement research via accurate and affordable smartphone eye tracking. Nature Communications, 11(1), Article 1. https://doi.org/10.1038/s41467-020-18360-5

Wallace, J. M., Chung, S. T. L., & Tjan, B. S. (2017). Object crowding in age-related macular degeneration. Journal of Vision, 17(1), 33. https://doi.org/10.1167/17.1.33

Watson, A. B., & Pelli, D. G. (1983). Quest: A Bayesian adaptive psychometric method. Perception & Psychophysics, 33(2), 113–120. https://doi.org/10.3758/BF03202828

Xia, R., & Barnes, G. (1999). Oculomanual Coordination in Tracking of Pseudorandom Target Motion Stimuli. Journal of Motor Behavior, 31(1), 21–38. https://doi.org/10.1080/00222899909601889

Xu, P., Ehinger, K. A., Zhang, Y., Finkelstein, A., Kulkarni, S. R., & Xiao, J. (2015). TurkerGaze: Crowdsourcing Saliency with Webcam based Eye Tracking (arXiv:1504.06755). arXiv. https://doi.org/10.48550/arXiv.1504.06755

