## Supplementary materials for "EasyEyes — Accurate fixation for online vision testing of crowding and beyond"

- Links to the EasyEyes github repository
  - https://github.com/EasyEyes/website (Website repo, ie easyeyes.app)
  - https://github.com/EasyEyes (EasyEyes org page, with all our repos)
  - https://github.com/EasyEyes/threshold (EE participant page, ie the actual experiment code)
- Link to the EasyEyes manual
  - https://docs.google.com/spreadsheets/d/1x65NjykMm-XUOz98Eu_oo6ON2xspm_h0Q0M2u6UGtug/edit#gid=1641458286
- Links to CriticalSpacing.m
  - https://github.com/denispelli/CriticalSpacing
- EasyEyesHandshaking.m
  - https://osf.io/u6gdj/
- Data
  - https://osf.io/u6gdj/
- Analysis and plotting code
  - https://osf.io/u6gdj/
- Kurzawski et al., 2023 on OSF
  - https://osf.io/83p6u/
- JS code for converting between coordinate systems
  - https://github.com/EasyEyes/threshold/blob/0a64d1257aaa2ef3a7b418fac766bcb6122a2eff/components/utils.js#L216 (XYPixOfXYDeg, deg (relative to fixation) to pix function. NOTE this function uses the psychoJS pixel coordinate system, wherein the origin is the center of the screen, increasing up and to the right. We convert this to the origin in top-left system that is standardized with MATLAB

using another function, linked below)

- https://github.com/EasyEyes/threshold/blob/0a64d1257aaa2ef3a7b418fac766bcb6122a2eff/components/utils.js#L277 (XYDegOfXYPix, pix to deg (relative to fixation) function. Again, this code assumes a coordinate system with the origin in the center of the screen as this is what PsychoJS methods expect.
- https://github.com/EasyEyes/threshold/blob/0a64d1257aaa2ef3a7b418fac766bcb6122a2eff/components/eyeTrackingFacilitation.ts#L134 (getAppleCoordinatePosition, function to convert between psychojs pixel coordinate system (ie origin at the center of the screen) to the Apple coordinate convention (ie origin at the top left), that is reported in the stimulus csv and used by MATLAB.
